# A male-specific neural circuit for appetitive thirst in Drosophila

**DOI:** 10.64898/2026.10.08.757749

**Authors:** Brian P. Wang, Xucheng Zhang, Sunaad Shastry, Gagandeep Kaur, Nailah Fatima, Fred W. Wolf

**Author notes:** Lead author: Fred W. Wolf.

## Abstract

Thirst is the brain representation of the need to restore water balance. While environmental and interoceptive sensory neurons that assess water availability and status are known, the central brain representation of thirst that determines behavior in Drosophila is less well understood. We identified a male-specific appetitive thirst circuit in Drosophila using circuit mapping, connectomics, and behavior measurements. Male-specific NPF-M neurons receive thirsty seeking information via Allatostatin A (AstA) neuropeptide released from the Janu-AstA projection neuron. In turn, NPF-M neurons release the neuropeptides sNPF, NPF, and DH31 to promote thirsty water seeking and the neurotransmitter acetylcholine to inhibit it. NPF-M neurons regulate multiple motivated behaviors, including appetitive and consummatory thirst, food foraging, and courtship, driven by distinct input neurons. NPF-M neurons output to appetitive and aversive dopaminergic, mushroom body ongoing behavior integration center, and navigation circuits. This circuit exhibits similarities to those for hungry food foraging and courtship. NPF-M and Janu-AstA form a network with OviIN hub neurons that are sex-independent thirst neurons. We suggest that NPF-M neurons form a circuit branch that funnels male-specific information to tune motivated behavior.

## Introduction

Thirst is the neurally represented drive for water seeking and ingestion elicited by high osmolar interstitial fluids and by low blood volume. Given the centrality of water homeostasis to life across the animal kingdom, robust neural circuits evolved to facilitate thirst-driven water seeking, water quality evaluation, ingestion, and the regulation of renal water retention ^1,2^. In the fly *Drosophila melanogaster*, water deprivation drives the same stereotyped actions ^3^. Flies afford the opportunity to define thirst representation at single neuron and whole circuitry level. Appetitive water seeking is a crucial motivational step that occurs on a timescale relevant to behavioral competition. Water seeking, like odor-tracking, is a continuous, sometimes long-distance process where new stimuli can refocus attention and risks may be encountered. Additionally, water seeking can facilitate other behaviors since sources of water are associated with food availability, conspecifics, and reproductive behaviors ^4–6^.

Circuits encoding thirst-motivated behavior lie deep in the brain, in higher order processing regions that are downstream of interoceptive and sensory neurons. Interoception of high osmolality interstitial fluids is detected by the interoceptive suboesophageal zone neurons (ISNs) that have neuronal cell bodies positioned outside the blood-brain barrier ^7,8^. Flies sense humidity via dedicated hygrosensory receptor neurons that confer humidity avoidance when water replete and humidity attraction when thirsty ^9–12^. Additional gustatory sensors detect water taste ^13,14^. The central brain circuitry for thirst begins to take shape with appetitive neurons downstream of interoceptive and sensory neurons. Notably, appetitive water seeking behavior is shaped by dopaminergic neurons in the PAM protocerebral anterior medial group, the αβ Kenyon cells of the mushroom body, the AstAergic projection neuron Janu-AstA, a GABAergic local interneuron in the suboesophageal zone, and neurons expressing the NPY-like Neuropeptide F (NPF) and its receptor NPFR ^15–18^. Water ingestion neural circuitry is also partly known, but is largely studied separately from appetitive thirst, increasing the challenge to distinguish thirst representation from step-specific circuitry ^13,14,19–23^. Finally, water reward in thirsty flies supports associative long-term memory formation through dopamine neurons, mushroom body learning and memory circuitry, and NPFR ^24–26^. Better understanding of the similar yet distinct circuits for appetitive, ingestive, and reward aspects of thirst representation will help reveal how complex motivated behavior is organized.

The anticipatory thirst neuron Janu-AstA is a useful entry point for understanding thirst representation. Whereas the state of thirst is negatively valanced, thirst-driven hygrotaxis and water intake are rewarding and positively valanced ^9,16,26–29^. Two components of the known thirsty water seeking circuitry encode positive valence, the PAM dopamine neurons and Janu-AstA, suggesting they may form part of an anticipatory reward pathway ^16,17,30^. Janu-AstA projects from the suboesophageal zone (SEZ) sensory processing brain region to the superior medial protocerebrum (SMP) higher order computational brain region, providing a pathway for sensory influence of ongoing behavior. Janu-AstA signals with the AstA neuropeptide to NPF neuropeptide-expressing neurons to regulate thirsty water seeking.

Here we leveraged Janu-AstA projection neuron connectivity and function as a starting point to discover thirsty water seeking neural circuitry. We unexpectedly uncovered sexual dimorphism in water seeking for the sexually isomorphic Janu-AstA. To identify the basis for this behavioral dimorphism, we identified and then determined the function of Janu-AstA output neurons (JAsONs) that are sexually dimorphic or that encode sex-specific behaviors.

## Results

### Thirsty water seeking is controlled sex-specifically by sexually isomorphic Janu-AstA neurons

We measured water seeking in water deprived (thirsty) males and females. To do this, we used an open field assay, where a group of 20 genetically identical flies (for n=1) are provided with a discrete inaccessible source of water in the center of the field ^16^. The water source sets up a 3-6% humidity gradient that flies can track. At 18 hours of water deprivation females showed less water seeking compared to males (**Figure 1A**). Extending the water deprivation time to 27 hours for females resulted in water seeking that is similar to males at 18 hours water deprivation. The sex difference in thirsty water seeking could be due to differences in body physiology, for example water balance in tissues, or in the response of the brain to thirst and humidity sensory cues. To test this, we used flies harboring mutations in the *fruitless* (*fru*) gene that selectively affect the sex of the brain ^31^. One mutation variant feminizes the brain but not the body of males (*fru^F^*), while the other mutation variant masculinizes the brain but not body of females (*fru^M^*). Males with feminized brains showed reduced water seeking, whereas females with a masculinized brain showed increased water seeking (**Figure 1B**). Thus, sex differences in thirsty water seeking are encoded by the brain.

**Figure 1.**
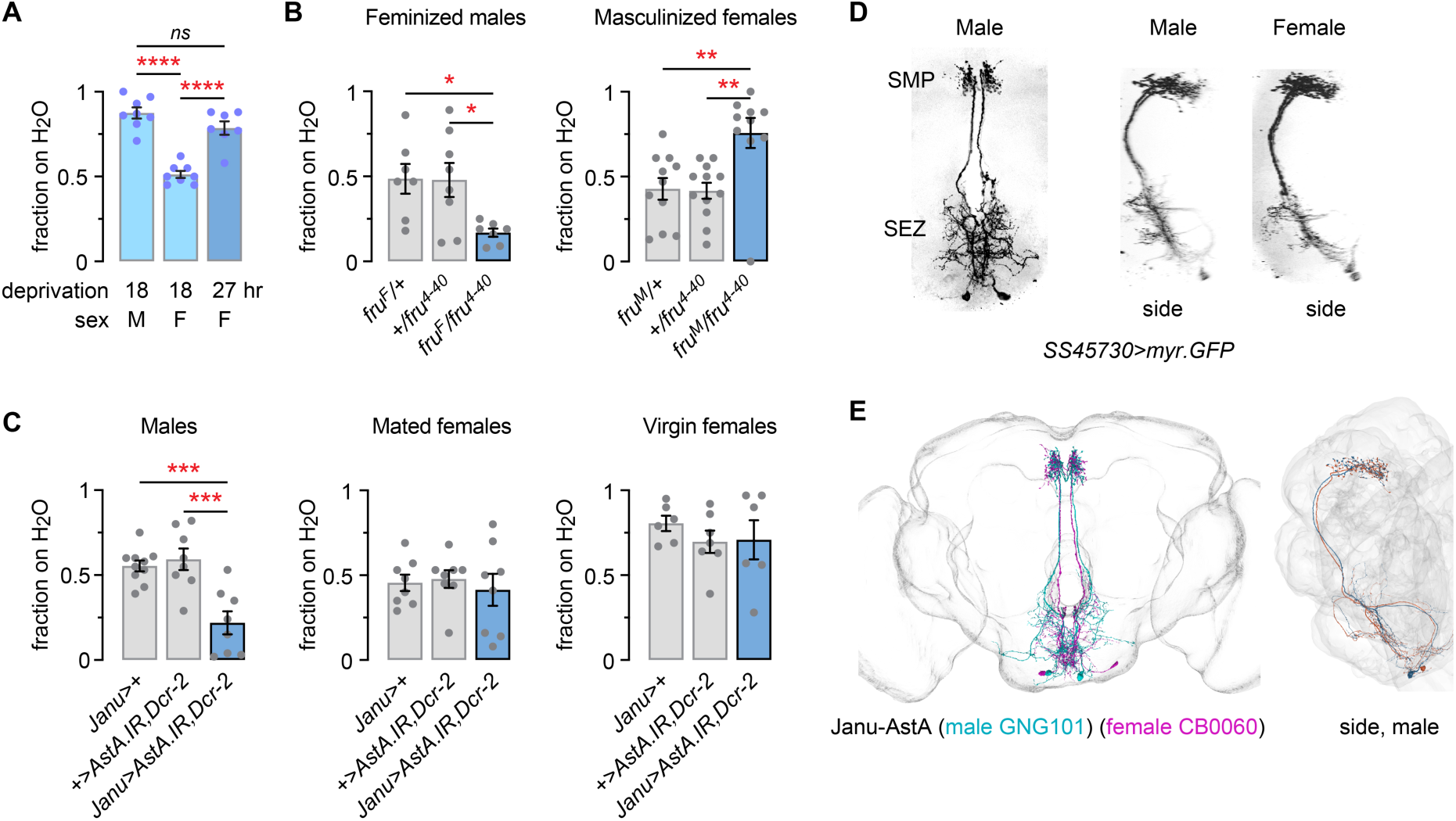
AstA in Janu-AstA is male-specific for sexually dimorphic thirsty water seeking. **A**. Fraction of male or female flies occupying a gridded water source following different lengths of water deprivation. One-way ANOVA/Tukey’s. **B**. Male flies with a feminized nervous system (*fru^F^/fru*^4–40^) exhibit female-like water seeking (left). Brown-Forsythe/Dunnett’s T3. Female flies with a masculinized nervous system (*fru^M^/fru*^4–40^) exhibit male-like thirsty water seeking (right). Kruskal-Wallis/Dunn’s. **C**. *Janu-spGal4* (*R52A01-AD;R65D05-DBD*) driving RNAi against AstA results in male-specific reductions in thirsty water seeking. Left: one-way ANOVA/Holm-Šídák’s; center: Brown-Forsythe; right: one-way ANOVA. **D**. No anatomical sexual dimorphism is revealed with Janu-AstA-specific *split-Gal4 SS45730*. Left: anterior view in male; right: side view in male and female. **E**. Left: Janu-AstA morphological comparison between the mCNS (GNG101) and FAFB (CB0060) connectomes. Right: side view in mCNS.

Brain encoding of the sex differences in water seeking led us to reexamine known water seeking circuitry. Janu-AstA is a projection neuron that carries positive valence water seeking drive from the gnathal ganglion (GNG) and flange suboesophageal zone (SEZ) neuropils to the superior medial protocerebrum (SMP) and releases the thirst neuropeptide AstA ^16^. Janu-AstA is the sole AstAergic neuron amongst the 4-5 neurons labeled by the Janu *split-Gal4* (*spGal4*, *R52A01-AD*;*R65D05-DBD*). Reduced AstA signaling in *Janu-spGal4* reduced water seeking in thirsty males, but it did not impact thirsty water seeking in either mated or virgin females (**Figure 1C**). We identified a second *spGal4*, *SS45730* (*R65D05-AD*;*VT027802-DBD*, formerly identified as labeling Trogon candidate feeding neurons), that specifically labels Janu-AstA ^32^. Janu-AstA neuronal morphology was not detectably different between males and females (**Figure 1D**). Neurons with the same unique morphology were identified in the Full Adult Female Brain (FAFB, CB0060) and the male CNS (mCNS, GNG101) connectomes, appearing sexually isomorphic (**Figure 1E**) ^33,34^. RNAi-mediated reduction of AstA with *SS45730-spGal4* also reduced water seeking in males but not in females (**Figure S1**). Thus, AstA release from Janu-AstA forms part of a male-specific water seeking circuit.

### Sexually dimorphic neural circuitry that is synaptically connected to Janu-AstA

We identified sexually dimorphic neurons that are synaptically connected to Janu-AstA, to ask if the male-specific function of Janu-AstA can be explained by its connectivity. First, we used the *trans*-Tango anterograde tracer to identify neurons postsynaptic to Janu-AstA in males ^35^. We colabeled for the *trans*-Tango reporter and the neuropeptide NPF, since NPF neurons function postsynpatic of Janu-AstA for water seeking ^16^. The anterograde label revealed dense output from Janu-AstA in the SMP, as expected (**Figure 2A**). Two groups of NPF-positive neurons were labeled: clock neurons of the circadian pacemaker and the male-specific NPF-M neurons. In the central clock, 4-5 of the 6 NPF-positive ventral group neurons, the lLNvs and the 5th LNv, were *trans*-Tango-positive, and 1-2 of the 3 NPF-positive dorsal group neurons, the LNd2/3 and LNd6, were *trans*-Tango-positive. Two NPF-M neurons were *trans*-Tango-positive. To characterize the NPF-Ms, we used *NPF-Gal4* combined with *fru-FLP* and *UAS-responder* transgenes that contain FRT-flanked translation stop cassettes ^36^. *fru-FLP* expresses Flippase in Fru-expressing neurons to excise the stop cassette and allow *UAS-responder* expression. *NPF-Gal4,fru-FLP>STOP.GFP* specifically labels 2-3 NPF-M neurons per hemisphere (**Figure 2B**). The morphology of a single NPF-M neuron was revealed using a stochastic labeling technique (**Figure 2C**) ^37^. The NPF-M bilaterally innervates the medial SMP region and the dorsal protocerebrum with presumed dendritic processes that were in close proximity to AstA release sites. The NPF-M neurons form an ipsilateral tilted ring of presynapses in the crepine brain region (**Figure 2D**). Closer inspection of the ring revealed a close association with the mushroom body horizontal lobes (**Figure 2E, F**). NPF-M synapses encircle the surface of the mushroom body Kenyon cells at the boundary of the γ3/γ4 compartments of the γ lobe. We identified two male-specific neurons in the mCNS connectome, SMP712m and ICL010m, as NPF-Ms (**Figure 2G**). SMP712m elaborates dendrites in the SMP and presynaptic release sites encircling the γ3/γ4 region of the mushroom body horizontal lobes, and ICL010m contributes presynaptic release sites to the dorsal aspect of the same presynaptic region as SMP712m. Thus, the male-specific NPF-Ms are post-synaptic to Janu-AstA, and they output information to the region surrounding the mushroom body horizontal lobe and to a lesser extent other parts of the dorsal protocerebrum.

**Figure 2.**
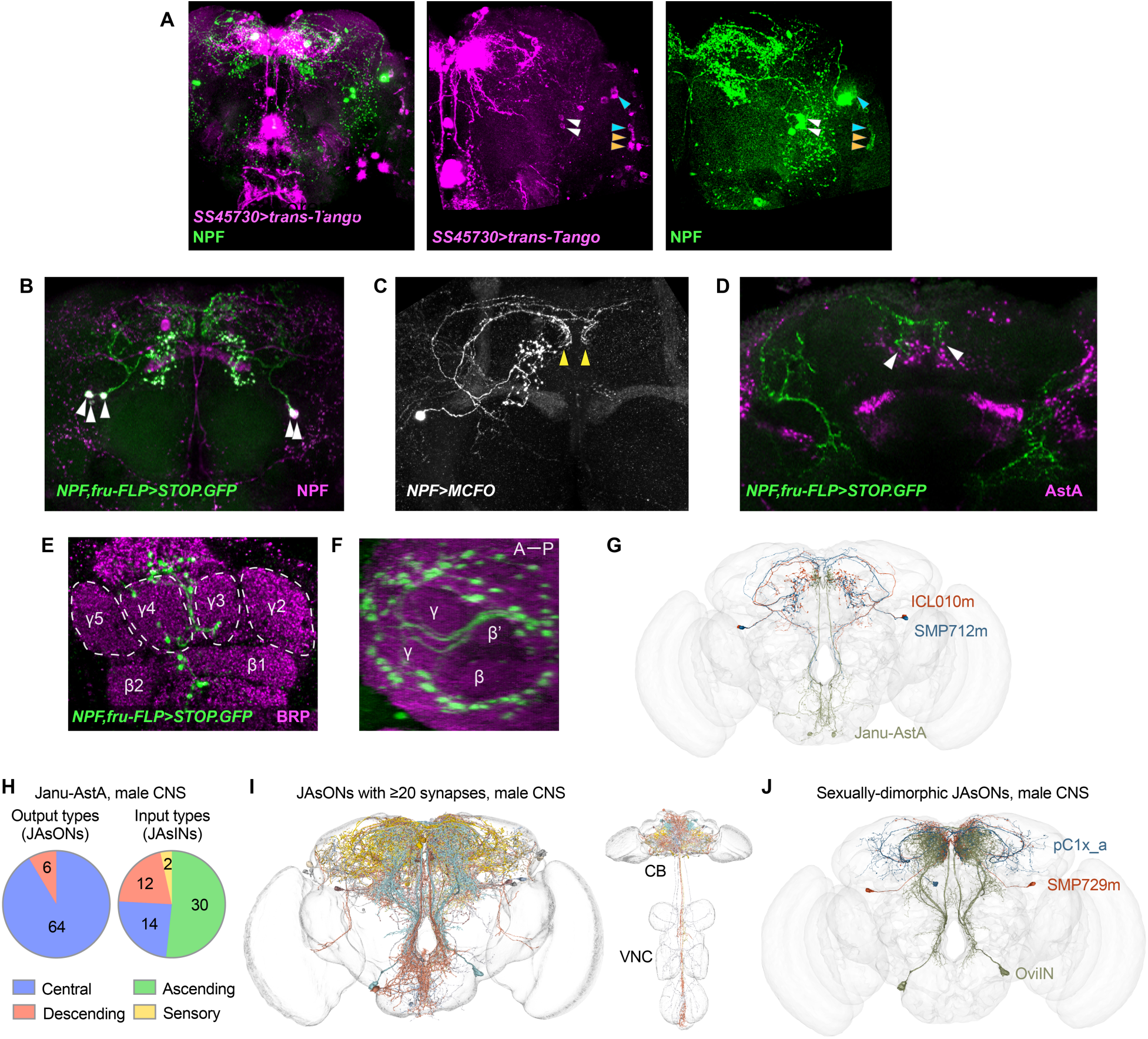
Janu-AstA synaptic output connectivity. **A**. *trans*-Tango anterograde synaptic tracing from Janu-AstA with *SS45730-spGal4* (magenta) and counterstained for NPF (green). White arrowheads point to NPF-positive NPF-M cell bodies, yellow arrowheads to NPF-positive ventral group circadian pacemaker neurons, and cyan arrowheads to NPF-positive dorsal group circadian neurons. **B**. Morphology of NPF-M neurons (green), counterstained for NPF (magenta). Arrowheads point to NPF-M cell bodies. **C**. A single SMP712m NPF-M neuron revealed by MultiColor FlpOut (MCFO). Arrowhead points to dendritic innervation of the SMP. **D**. Apposition of NPF-M dendrites and AstA immunostaining in the SMP (arrowheads). **E**. NPF-M presynaptic release sites (green) at the surface of the γ3 and γ4 compartments of the mushroom bodies. Counterstained with the BRP presynaptic marker (magenta), single 1 μm confocal section. **F**. The same region as in (E), rotated horizontally by 90°, revealing the ring of NPF-M presynapses surrounding the β, β’, and γ mushroom body lobes. **G**. NPF-M neurons SMP712m and ICL010m and Janu-AstA (GNG101) in the male CNS connectome. **H**. Janu-AstA output (JAsON) and input (JAsIN) neuron types. Values represent all neuron types making 5 or more synapses with Janu-AstA. **I**. JAsONs making 20 or more synapses with Janu-AstA, in the mCNS. **J**. Sexually dimorphic (pC1x_a and OviIN) and sex-specific (SMP729m) JAsONs from the 20 or more synapses with Janu-AstA cutoff in Figure 2I.

We categorized the synaptic partners of Janu-AstA in the mCNS connectome (**Figure 2H**) ^33^. Janu-AstA output neurons, or JAsONs, are predominantly comprised of central brain neurons, with the remainder being descending neurons that connect the central brain to the VNC. Janu-AstA sends output to 95 (left) and 98 (right) neurons, or 83 unique neurons after removing contralateral homologs, that are comprised of 70 neuron types (**Figure 2I, Table S1**). The connectome did not reveal synaptic connections between Janu-AstA and NPF-M neurons or NPF-positive clock neurons. The reconstruction of fine dendritic processes is incomplete in the mCNS and the other extant connectomes ^33^. Additionally, synaptic connectivity can vary based on prior experience, physiological state, and between individuals. In the mCNS, Janu-AstA neuronal processes in the SMP are 0-0.31 μm from NPF-M SMP712m and 0.93-23.2 μm from NPF-M ICL010m. Finally, three of 19 highly connected JAsONs (>50 synapses per neuron type and more than 10 synapses per individual neuron within type) are sexually dimorphic or male specific: Oviposition inhibitory neuron (OviIN), pC1x_a, and SMP729m (**Figure 2J**). These neurons are further characterized below, in the context of the roles of the NPF-M and the OviIN in thirst.

Janu-AstA receives synaptic input from 74 (left) and 82 (right) neurons, or 75 unique neurons after removing contralateral homologs, that are comprised of 58 neuron types (**Figure 2H**). Half of Janu-AstA inputs, or JAsINs, are ascending neurons (ANs) and sensory neurons, with synaptic outputs in the central brain and cell bodies in the VNC and the periphery. Other Janu-AstA synaptic partners include descending neurons (DNs) and SEZ interneurons. Direct sensory input to Janu-AstA is predominantly from 6 of 12 total SAxx02 neurons. SAxx02 is a group of leg sensory neurons of an unknown sensory modality ^38^. We also assessed the 11 ANs that make 50 or more synapses with Janu-AstA for second order sensory input that arises from the thorax and abdomen (**Figure S2A**). SAxx02, in addition to its direct input to Janu-AstA, was presynaptic to seven of eleven JAsIN ANs, making a total of 6,197 synaptic connections or 27% of SAxx02 output. Seven other sensory neuron types (SNch01, SNch10, SNpp32, SNta04, SNta18, SNxx25, and SNxx29) provide input to three or more JAsINs. These sensory types transduce chemosensory (ch), proprioceptive (pp), tactile (ta), and unknown (xx) information. SNch01 provides the largest number of synaptic inputs to a JAsIN, making 12,923 synapses specifically with the ascending JAsIN AN09B018. AN09B018 itself receives strong multimodal sensory information, including from SAxx02 and SNxx29. Thus, Janu-AstA receives direct sensory information from each leg via SAxx02 and indirect sensory information from an array of body sensors. We sought to determine whether hygrosensory information constitutes a portion of the sensory input to Janu-AstA. We did not detect Ir8a, Ir40a, or Ir68a hygrosensory neuron innervation of the legs or wings (**Figure S2B**).

Two other features of Janu-AstA input may be relevant to thirsty seeking. First, DNp48 is the sole sexually dimorphic JAsIN ^39^. DNp48 connectivity and possible function are discussed in the context of the NPF-M. Second, the GNG neuron GNG495 and Janu-AstA uniquely share a pattern of synaptic inputs (**Figure S2C**). Both neurons are highly interconnected with AN09B018, AN05B096, and AN05B100. GNG495 is morphologically similar to Janu-AstA in the SEZ, but it differs in that its inputs and outputs largely confined to the SEZ. Hence, Janu-AstA and GNG495 may be coregulated by overlapping inputs to provide output to the SMP and the SEZ, respectively.

### NPF-M neurons regulate water seeking in a thirst-dependent manner via multiple neuropeptides and acetylcholine

Because NPF-M neurons may provide male-specificity to the Janu-AstA water seeking circuit, we tested their role in thirsty water seeking. Blocking presynaptic release in NPF-M neurons with tetanus toxin light chain (TeTx) increased thirsty water seeking in males, but not in females as expected, since NPF-Ms are male-specific (**Figure 3A**). We tested for an effect in fully water replete males, and we observed no increase in water seeking (**Figure 3B**). Thus, synaptic release from NPF-M neurons inhibits thirsty water seeking specifically in thirsty flies. This thirst dependence for NPF-M function suggests that NPF-M neurons modify the function of central thirst circuits. NPF-M acute activation using the heat activated TrpA1 cation channel decreased thirsty water seeking in males (**Figure 3C**). Thus, ongoing NPF-M neuronal activity can bidirectionally regulate thirsty water seeking in male flies.

**Figure 3.**
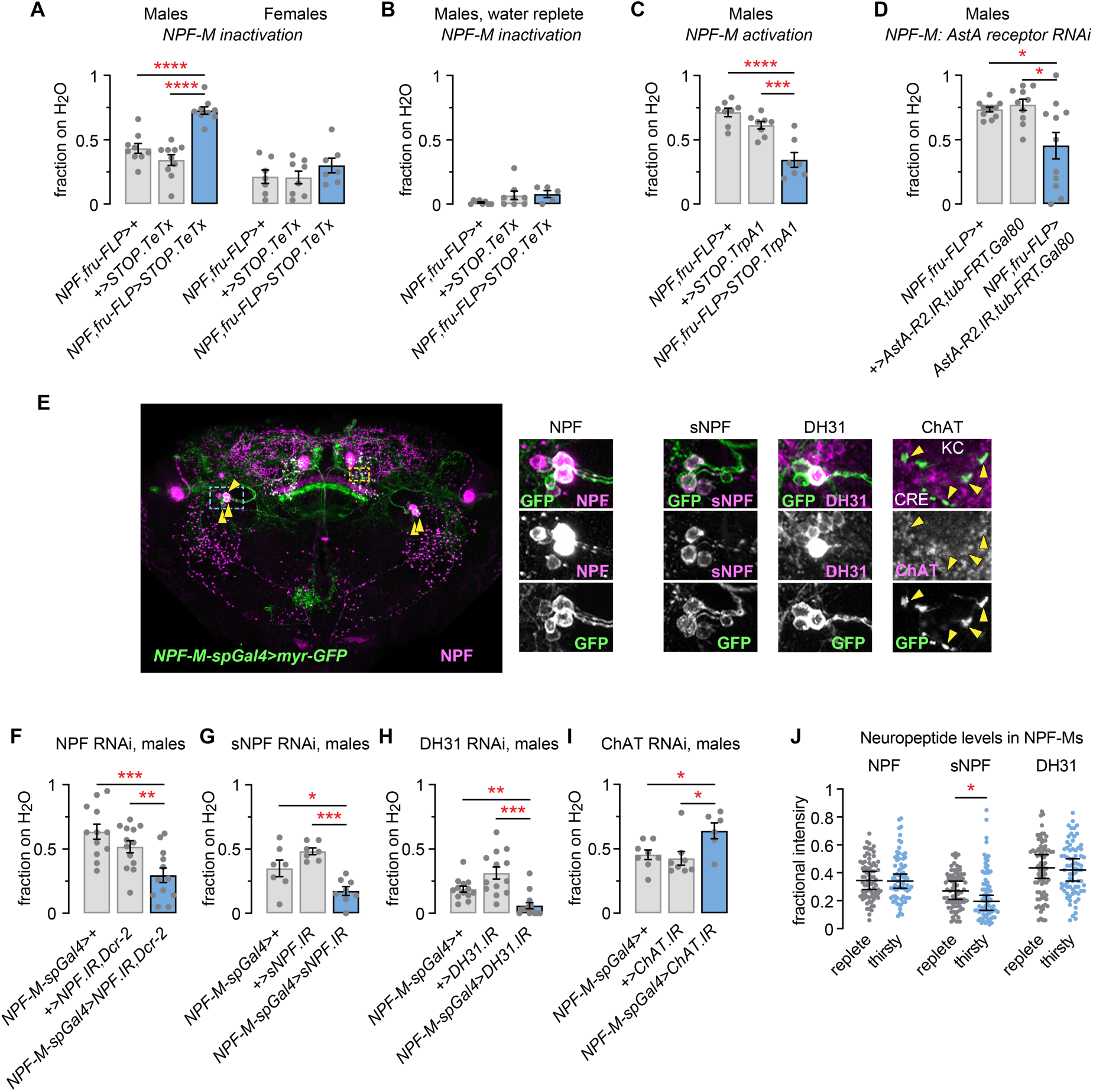
Cotransmission of three neuropeptides and acetylcholine by NPF-M neurons shapes thirsty water seeking. **A**. Inactivation of NPF-M neurons with the tetanus toxin light chain (TeTx) increases thirsty water seeking in males but not females. Left: one-way ANOVA/Holm-Šídák’s; right: one-way ANOVA/Holm-Šídák’s. **B**. No effect of inactivation of NPF-M neurons on water seeking in water replete males. Kruskal-Wallis. **C**. Acute activation of NPF-Ms with TrpA1 decreases thirsty water seeking. One-way ANOVA/Holm-Šídák’s. **D**. Decreased expression of the AstA-R2 receptor in NPF-M neurons decreases thirsty water seeking. In the experimental genotype (*NPF-Gal4;fru-FLP>AstA-R2.IR;tub-FRT.Gal80*), the GAL4 repressor, GAL80, is expressed in all cells except for NPF-M neurons. Brown-Forsythe/Dunnett’s T3. **E**. *NPF-M-spGal4* expression pattern, counterstained for NPF (left panel). Panels to the right are of the cell body region indicated for the blue dashed box, stained for GFP and either NPF, sNPF, or DH31. Right-most panel is the yellow dashed box NPF-M presynaptic region, stained for ChAT. KC: Kenyon cell, CRE: crepine brain region. **F**. RNAi against NPF in NPF-M neurons reduces thirsty water seeking. One-way ANOVA/Holm-Šídák’s. **G**. RNAi against sNPF in NPF-M neurons reduces thirsty water seeking. One-way ANOVA/Holm-Šídák’s. **H**. RNAi against DH31 in NPF-M neurons reduces thirsty water seeking. Brown-Forsythe/Dunnett’s T3. **I**. RNAi against ChAT in NPF-M neurons increases thirsty water seeking. One-way ANOVA/Holm-Šídák’s. **J**. Neuropeptide levels in NPF-M cell bodies for NPF, sNPF, and DH31. Data is represented as fractional mean pixel intensity for individual cell bodies. NPF: Two-tailed t test; sNPF: two-tailed Mann-Whitney test; DH31: two-tailed Welch’s t test.

To test if NPF-M neurons can receive an AstAergic signal to regulate water seeking, we used an additional FRT recombination tool, *tub-FRT.Gal80*, where the GAL4 inhibitor, GAL80, is selectively excised in FLP-expressing cells. This allows expression of *UAS-responder* transgenes in NPF-M neurons with the *NPF-Gal4;fru-FLP* NPF-M driver. In flies of the genotype *NPF-Gal4,fru-FLP,tub-FRT.Gal80,UAS-AstA-R2.IR*, RNAi against the AstA receptor, AstA-R2, is expressed specifically in NPF-M neurons. *AstA-R2* was required in NPF-M neurons to promote thirsty water seeking, strengthening the likelihood that AstA release from Janu-AstA signals to NPF-M neurons to regulate thirsty water seeking (**Figure 3D**).

To allow broader testing of *UAS-responder* transgenes and the function of NPF-Ms, we created a new *split-Gal4*, *NPF-M-spGal4* (*VT002860-AD;NPF-T2A-DBD*), that expresses in 2-3 NPF-M neurons (**Figure 3E**). *NPF-M-spGal4* also expresses in the NPF-positive small DM neurons and in NPF-negative neurons in the fan-shaped body. *NPF-M-spGal4* is not expressed in any intestinal cells including NPF-positive enteroendocrine cells of the intestine that are implicated in regulating nutritional physiology (**Figure S3A**) ^40,41^. To test if *NPF-M-spGal4* and *NPF-Gal4,fru-FLP* are similar for the purpose of manipulating NPF-Ms, we sought to recapitulate the effects of *NPF-Gal4,fru-FLP* with *NPF-M-spGal4.* Expression of TeTx to block synaptic transmission in *NPF-M-spGal4* increased thirsty water seeking in males (**Figure S3B**). *AstA-R2* RNAi in *NPF-M-spGal4* decreased thirsty water seeking (**Figure S3C**). Thus, *NPF-M-spGal4* and *NPF-Gal4,fru-FLP* neuron manipulation result in concordant behavioral phenotypes.

We next identified neuromodulators and neurotransmitters released by NPF-M neurons. We suspected that NPF-M neurons may co-release neuromodulators or neurotransmitters in addition to NPF, since blocking NPF-M synaptic release increased thirsty water seeking, the opposite effect from reducing NPF expression in all NPF neurons ^16^. Previously published single cell sequencing data was used to identify other neuromodulators and neurotransmitters in NPF-Ms. Whole head single cell sequencing data was gated for the presence of neuronal markers (*elav*, *nSyb*), the central nervous system (*oc*), *NPF*, and *fru* ^42^. We also performed a simple lookup on *fru*^+^ single cell sequencing data tables derived from male pupae ^43^. The selected putative NPF-M cells were positive in one or both datasets for the neuropeptides NPF, sNPF, DH31, and CCHa2. Putative NPF-M neurons were also positive for the vesicular acetylcholine transporter (VAChT) and the Ddc serotonergic biosynthetic enzyme. Immunohistochemistry revealed that NPF-M neurons co-expressed the neuropeptides sNPF, NPF, and DH31, and the acetylcholine biosynthetic enzyme choline acetyltransferase (ChAT) (**Figure 3E**). We used RNAi to test the role of NPF, sNPF, DH31, and acetylcholine in NPF-M neurons. RNAi-mediated reduction of NPF, sNPF, or DH31 in *NPF-M-spGal4* each reduced thirsty water seeking in males but not in females (**Figure 3F-H, Figure S3D-F**). Thus, three neuropeptides function in NPF-M neurons to promote thirsty water seeking in males. Moreover, NPF-M acetylcholine acts oppositely to limit thirsty water seeking (**Figure 3I**). Acetylcholine release may thus explain the effects of manipulating the acute neuronal activity of the NPF-M neurons (**Figure 3A**, **C**). We asked if water deprivation affects neuropeptide levels NPF-M neurons. sNPF, but not NPF nor DH31, showed reduced intensity cell body staining in thirsty males, suggesting that thirst causes release of sNPF by the NPF-Ms (**Figure 3J**). The rich repertoire of signaling molecules required by the NPF-M neurons suggests that NPF-M neurons simultaneously release signals to create multifaceted postsynaptic responses.

### NPF-M neurons synapse onto neurons for memory, navigation, and sexually dimorphic behaviors

To ask how NPF-M neurons might function in a circuit to regulate thirst, we determined synaptic connectivity of SMP712m and ICL010m in the mCNS. We focused on neuron types that make 30 or more synapses with NPF-M (**Figure 4A**). SMP712m outputs are dominated by the PAM dopamine neuron (DAN) type PAM08 (20 of 50 PAM08 neurons, receiving 19% or 224 synapses of SMP712m output) with a smaller contribution (3%) of the PAM07 neurons (**Figure 4B**). PAM08 and PAM07 are also postsynaptic to ICL010m. PAM08 (PAM-γ4) and PAM07 (PAM-γ4<γ2γ1) (collectively PAM-γ4) innervate the γ4 compartment of the mushroom body horizontal lobes ^30^. PAM-γ4 DANs mediate appetitive learning and memory, including thirst-dependent water reward memory ^17,26,44^. PAM-γ4 dendrites receive synaptic input from NPF-M neurons just external to the mushroom body intrinsic Kenyon cells. SMP712m output neurons also include the fan-shaped body (FB) neuron types FB4G (4% of NPF-M output), FC2B (4%), FB4K (3%), and FC2A (3%) for a total of 14% of SMP712m output. The FB is part of the central complex that is important for orientation and navigation as well as other behaviors ^45^. ICL010m is presynaptic to the FB neuron type FB4G (2.6% of ICL010m output), and ICL010m is an NPF-M output to the OviIN.

**Figure 4.**
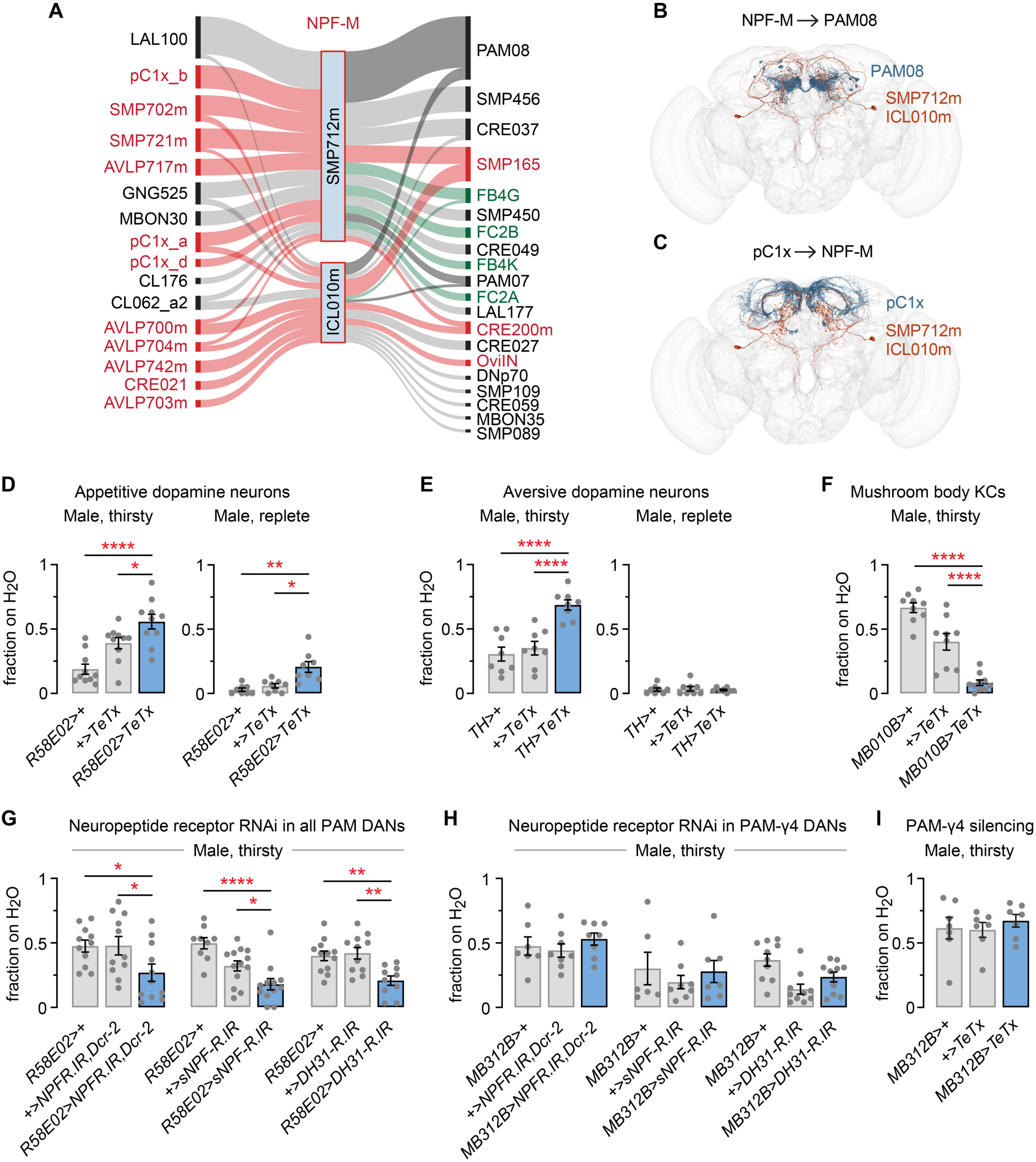
NPF-M output to dopaminergic neurons and mushroom body Kenyon cell (KC) neurons for thirsty water seeking. **A**. NPF-M input (left) and output (right) neuron types that make 30 or more synapses with NPF-M. Red color indicates sexually dimorphic or sex-specific neurons, and green indicates fan-shaped body neuron types. **B**. Morphology of NPF-Ms and the PAM08 PAM-γ4 neuron type. **C**. Morphology of the pC1x (pC1x_a, pC1x_b and pC1x_d) neurons that are inputs to NPF-M neurons. **D**. Synaptic inactivation of PAM dopamine neurons increases water seeking in both thirsty and water replete flies. Left: one-way ANOVA/Holm-Šídák’s; right: Brown-Forsythe/Dunnett’s T3. **E**. Synaptic inactivation of aversive dopamine neurons increases water seeking specifically in thirsty flies. Left: one-way ANOVA/Holm-Šídák’s; right: Kruskal-Wallis test. **F**. Synaptic inactivation of all mushroom body KCs decreases water seeking in thirsty flies. One-way ANOVA/Holm-Šídák’s. **G**. RNAi against the NPF receptor NPFR, the sNPF receptor sNPF-R, and the DH31 receptor DH31-R in all PAM dopamine neurons deceases thirsty water seeking. One-way ANOVA/Holm-Šídák’s. **H**. RNAi against the NPFR, sNPF-R, and DH31-R in the PAM-γ4 dopamine neurons has no effect on thirsty water seeking. One-way ANOVA. **I**. Synaptic inactivation of the PAM-γ4 neurons has no effect on thirsty water seeking. One-way ANOVA.

Sixteen neuron types provide 30 or more input synapses to NPF-Ms (**Figure 4A**). Eleven of these inputs (69%) are sexually dimorphic or sex specific. The sexually dimorphic pC1x neurons (12.4% input to both NPF-Ms, comprising 17% of SMP712m input and 5% of ICL010m input), including pC1x_b (9% of SMP712m input), pC1x_a (5% of each NPF-M), and pC1x_d (3% of SMP712m input), regulate male courtship and aggression (**Figure 4C**) ^46,47^. The NPF-M neurons also receive male-specific inputs from neurons innervating the anterior ventrolateral protocerebrum (AVLP) that are of unknown function. By comparison, only three of 20 (15%) NPF-M output neuron types are sexually dimorphic. The sexually dimorphic input bias suggests that a function of NPF-M neurons is to funnel sex-specific information into sexually isomorphic PAM and FB circuitry to influence the function of sex-independent circuits.

### Neuropeptide regulation of dopamine and mushroom body circuitry for thirsty water seeking

NPF-M synaptic output to dopaminergic neurons and the role of mushroom body circuitry in humidity attraction and water reward memories led us to test this circuitry for a role in thirsty water seeking. First, we blocked synaptic release in appetitive dopamine neurons (*R58E02-Gal4*), aversive dopamine neurons (*TH-Gal4*), and intrinsic mushroom body Kenyon cells (*MB010B-Gal4*). Blocking synaptic release in appetitive dopamine neurons increased thirsty seeking in both thirsty and water replete males (**Figure 4D**). The same manipulation in aversive dopamine neurons caused a thirst-specific increase in water seeking in males (**Figure 4E**). Thus, both appetitive and aversive dopamine neurons moderate thirsty water seeking. Expression of TeTx in the Kenyon cells (KC) with the KC-specific driver MB010B reduced thirsty water seeking, in agreement with prior findings (**Figure 4F**) ^15^. Interestingly, synaptic silencing with the widely used pan-KC driver *R13F02-Gal4* caused dramatic thirst-specific lethality (**Figure S4A**). *R13F02-Gal4* is expressed in a small number of neurons other than the KCs (**Figure S4B**). *R13F02>TeTx* flies given a shorter 4 hr water deprivation showed decreased water seeking, the same as with *MB010B>TeTx* (**Figure S4C**). Thus, KCs promote thirsty water seeking, and non-KC neurons in *R13F02-Gal4* ensure viability when flies are thirsty. In summary, aversive dopamine neurons, appetitive dopamine neurons, and mushroom body KCs all contribute to thirsty water seeking.

NPF-M neuropeptides might signal to mushroom body circuitry for thirsty water seeking. Expression of RNAi against any of the three receptors for the NPF-M neuropeptides, NPFR, sNPF-R, and DH31-R, in appetitive PAM dopamine neurons reduced thirsty water seeking in males (**Figure 4G**). Consistent with sNPF release from NPF-M acting on appetitive dopamine neurons, RNAi against sNPF-R in these neurons in females had no effect (**Figure S4D**). We also performed a partial assessment of neuropeptide receptor function in aversive dopaminergic neurons (*NPFR*) and the KCs (*sNPF-R*), and we found that both were required to promote thirsty water seeking (**Figure S4E, F**). Thus, it appears that NPF, sNPF, and potentially DH31 act broadly in dopaminergic and mushroom body KCs to regulate thirsty water seeking. Because NPF-M neurons are synaptically connected to the PAM-γ4 appetitive dopamine neurons, we directly tested the role of these neurons using the *MB312B-spGal4* driver. RNAi against *NPFR*, *sNPF-R*, or *DH31-R* in the PAM-γ4 neurons had no effect on thirsty water seeking (**Figure 4H**). A second, independent RNAi against *sNPF-R* also had no effect when expressed in the PAM-γ4 neurons (**Figure S4G**). Since NPF-M neuropeptides did not act through the PAM-γ4 neurons, we asked if PAM-γ4 neurons might instead be coupled to NPF-M cholinergic neurotransmission. Synaptic inactivation the PAM-γ4 neurons had no effect on thirsty water seeking (**Figure 4I**). This result partially segregates the mushroom body circuitry for thirst-driven innate water seeking and water reward learning. Other, currently unidentified neurons inhibit thirsty water seeking via cholinergic signaling from NPF-M neurons.

### Functional role of NPF-M circuitry

We broadened NPF-M characterization to ingestive behaviors to better define their core function. We tested thirsty water intake to determine if NPF-M neurons more generally represent the state of thirst. Blocking NPF-M synaptic release or decreasing expression of NPF in NPF-Ms increased total time of water intake, with no change in cibarial pumping frequency, indicating increased water intake (**Figure 5A**, **B**). Thus, NPF-M neurons regulate both the appetitive and consummatory phases of thirst. NPF-M neurons could specifically regulate thirst, or they could function across ingestive behaviors. To test this, we made flies hungry but not thirsty (wet starved), and tested foraging behavior towards a discrete source of dry food ^48^. Blocking synaptic release from NPF-Ms increased foraging for food (**Figure 5C**). Thus, NPF-M neurons appear to represent an aspect of internal need state that is common between thirst, hunger, and male mating drive.

**Figure 5.**
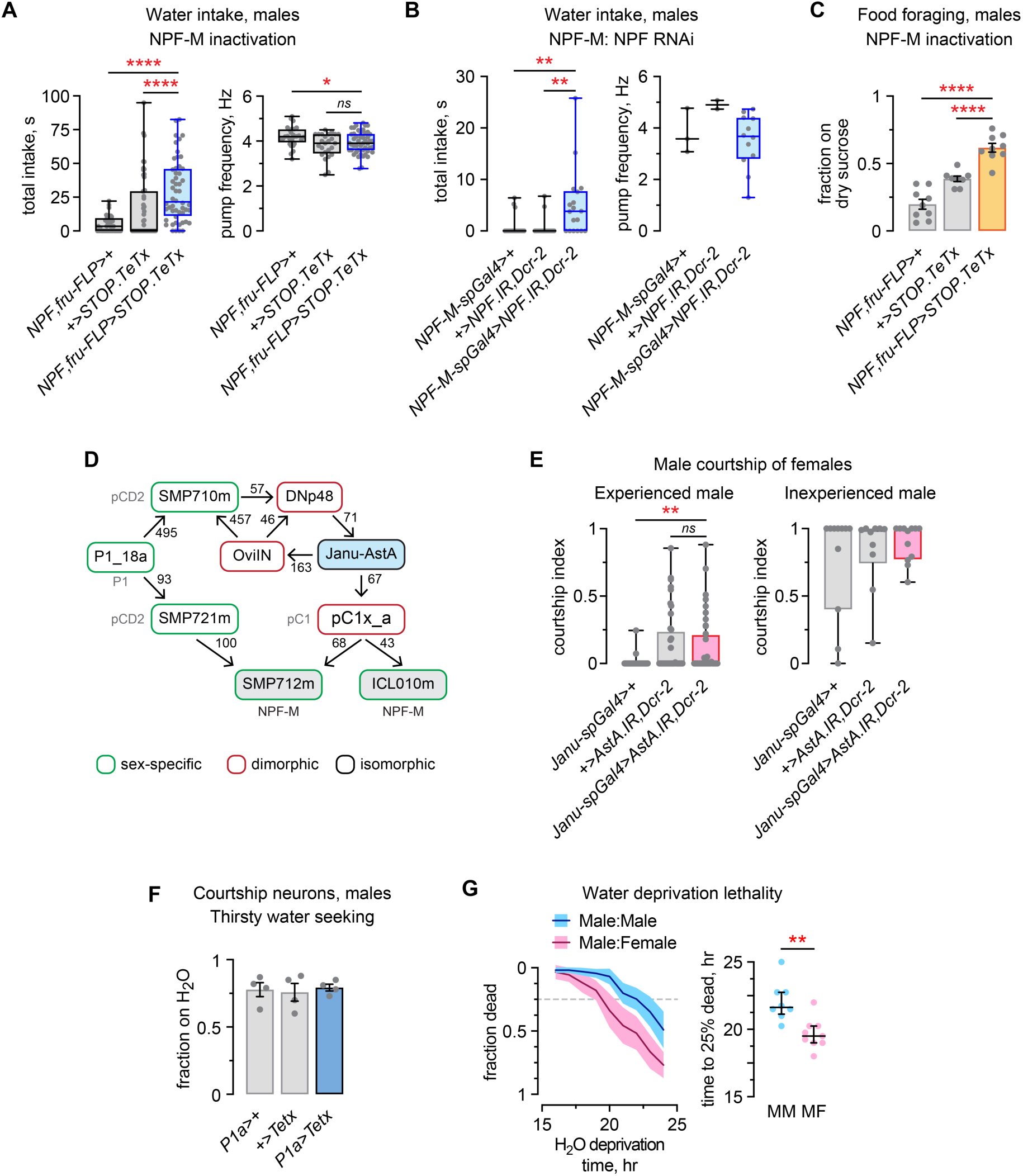
NPF-M neurons inhibit multiple male behaviors and function in a state-dependent manner. **A**. Increased water intake with inactivation of NPF-M neurons, total intake time (left) and cibarial pumping frequency (right). Left: Kruskal-Wallis test/Dunn’s; right: one-way ANOVA/Holm-Šídák’s. **B**. Increased water intake with reduced NPF expression in NPF-M neurons. Left: Kruskal-Wallis test/Dunn’s; right: one-way ANOVA. **C**. Increased foraging for food in open field assay with presentation of a discrete source of dry sucrose to wet starved flies. One-way ANOVA/Holm-Šídák’s. **D**. Sex-specific and sexually dimorphic neurons connecting NPF-M neurons and Janu-AstA. **E**. Experienced (left) and inexperienced (right) male courtship of females in flies with reduced AstA signaling in Janu-AstA neurons. Left: Kruskal-Wallis test/Dunn’s; right: Kruskal-Wallis test. **F**. Inactivating P1a courtship neurons does not affect thirsty seeking. One-way ANOVA. **G**. Sensitivity to water deprivation of sex-deprived males cohoused with either other males or mated females. Unpaired two-tailed t test.

The male-specificity of Janu-AstA/NPF-M thirst circuit and the role of NPF-M neurons in male courtship suggests an interaction between thirst and sexual behavior ^36,49^. Alternatively, NPF-M neurons may perform a more fundamental function that is common to a variety of otherwise distinct motivated behaviors. We first asked how sexually dimorphic neural circuitry integrates with NPF-M and Janu-AstA in the male connectome (**Figure 5D**). The P1_18a neuron is a member of the P1 group of male courtship neurons and that is unique amongst P1 neurons in receiving inputs from a broad array of P1 neurons ^47^. P1_18a is presynaptic to the male-specific SMP710m and SMP721m pCD2 neurons that may encode persistence in social interactions ^50^. The SMP721m pCD2 neuron links P1_18a to the SMP712m NPF-M neuron. The second pCD2 neuron, SMP710m, is upstream to Janu-AstA through the sexually dimorphic DNp48. Hence, male-specific information may be transmitted to Janu-AstA through this network. Moreover, the GABAergic and sexually dimorphic OviIN provides a potential feedback mechanism from Janu-AstA back to SMP710m and DNp48 to inhibit the circuit. Janu-AstA may transmit DNp48-processed information to NPF-M neurons through the pC1 neuron, pC1x_a. Thus, activation of P1_18a could lead to co-regulation of Janu-AstA and NPF-M and contribute to male-specific behavior.

We tested whether NPF-M presynaptic neurons for thirst behavior, Janu-AstA, and sexual behavior, P1a, share behavioral functions. We first asked if Janu-AstA is part of the male courtship circuitry. RNAi reduction of AstA in Janu-AstA did not increase male courtship of females by sexually experienced males, nor did it reduce courtship by sexually inexperienced males (**Figure 5E**). P1a neurons are courtship and aggression promoting neurons that act upstream of NPF-M neurons ^36^. Using the P1a *spGal4* driver *R15A01-AD;R71G01-DBD,* we silenced the neurons and observed no effect on male thirsty water seeking (**Figure 5F**). These findings suggest that thirst and courtship signals converge onto NPF-M neurons through separate pathways, and that NPF-M neurons perform a function for the predominant internal state. Finally, we tested for a behavioral relationship between thirst and courtship. Sex-deprived male flies were water deprived either in the presence or absence of females, to test the effect of courtship opportunity on thirst. The presence of females increased male sensitivity to water deprivation (**Figure 5G**). Thus, courtship increases thirst in males.

### Central brain neurons that regulate thirsty water seeking in both males and females

The thirsty water seeking neural circuit we uncovered does not explain sex-generalized thirst drive. Thus, we asked if other neurons postsynaptic to Janu-AstA or central brain neurons for water ingestion might function in thirsty water seeking in both sexes. The GABAergic OviIN accepts the 9th most synaptic output from Janu-AstA (163 of 4409 total Janu-AstA synapses in mCNS), inhibits female oviposition, and is sexually dimorphic (**Figure 6A**) ^51^. To test the role of the OviIN in thirsty water seeking, we used *OviIN-SS2 (VT026347-AD;VT026035-DBD)*, a previously characterized *split-Gal4* driver, and we created *OviIN-SS3* (*R68A10-AD;VT026035-DBD*) that has increased OviIN specificity (**Figure 6B** and **Figure S5A**). Reducing GABAergic signaling in the OviIN by RNAi against the GABA biosynthetic enzyme, Glutamic acid decarboxylase 1 (Gad1), with either *OviIN-SS2* or *OviIN-SS3* decreased thirsty water seeking in both males and in females (**Figure 6C** and **S5B**). The OviIN receives an AstAergic signal via the AstA-R2 receptor to regulate thirsty water seeking (**Figure 6D**). Thus, the OviIN promotes thirsty water seeking in both males and females. These results raised the possibility that Janu-AstA may also function in oviposition and that oviposition may be related to water seeking. Reduction of AstA expression in Janu-AstA failed to alter egg laying in either mated females or virgin females (**Figure 6E**). Since Janu-AstA is male specific for thirst, the state of thirst in females may regulate the OviIN through a separate circuit.

**Figure 6.**
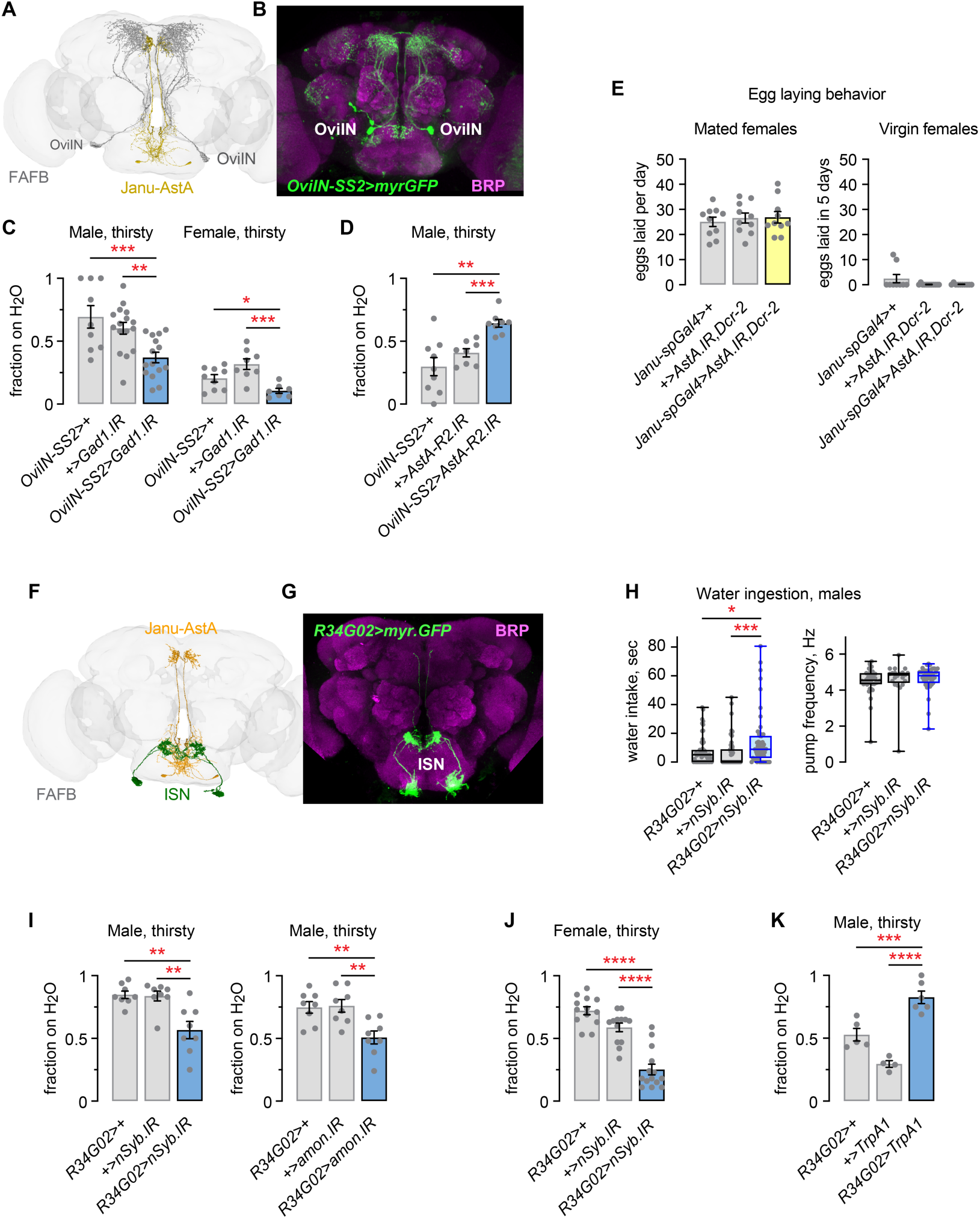
Neurons that regulate thirsty water seeking in both males and females. **A**. FAFB reconstruction of the OviIN neurons and the Janu-AstA neurons. **B**. Expression pattern of the *OviIN-SS2 split-Gal4*. **C**. RNAi reduction of Gad1 with OviIN*-SS2* reduces thirsty water seeking in males and females. Left and right: one-way ANOVA/Holm-Šídák’s. **D**. RNAi reduction of AstA-R2 in *OviIN-SS2* increases thirsty water seeking in males. Brown-Forsythe/Dunnett’s T3. **E**. RNAi reduction of AstA with *Janu-spGal4* has no effect on egg laying in either mated or virgin females. Left: one-way ANOVA; right: Kruskal-Wallis. **F**. ISN output near Janu-AstA dendrites in the FAFB. **G**. *R34G02-Gal4* is expressed specifically in the ISNs. **H**. Increased water intake with neural synaptobrevin (nSyb) RNAi that blocks synaptic release in the ISNs, total intake time (left) and cibarial pumping frequency (right). Left: Kruskal-Wallis/Dunn’s; right: Kruskal-Wallis. **I**. RNAi against nSyb in the ISNs decreases thirsty water seeking in males (left). RNAi against amontillado (amon) in the ISNs decreases thirsty water seeking in males (right). Left and right: one-way ANOVA/Holm-Šídák’s. **J**. nSyb RNAi in the ISNs decreases thirsty water seeking in females. One-way ANOVA/Holm-Šídák’s. **K**. Activation of the ISNs with the heat-activated cation channel TrpA1 increased thirsty water seeking in males. One-way ANOVA/Holm-Šídák’s.

The ISNs are candidates for initiating the state of thirst. High osmolarity inhibits the hemolymph-exposed ISNs to promote water intake in females ^8,19,52^. It remains unknown whether the ISNs regulate thirst in males and whether they coregulate both appetitive and consummatory thirst behavior. The ISNs make a dense innervation into a confined area of the prow and flange brain regions, in close proximity (0.03-1.13 μm in mCNS) to Janu-AstA dendrites (**Figure 6F**, **G**). Additionally, the GNG323 projection neuron from the SEZ to the SMP may provide a relevant functional connection from the ISN (29 synapses) to the OviIN (216 synapses). We inhibited the ISNs (*R34G02-Gal4*) using genetic conditions previously reported to impact water intake: decreasing the expression of neural Synaptobrevin (nSyb) to block neurotransmission, decreasing expression of the Amontillado (Amon) peptidase that cleaves the pro-forms of neuropeptides, and expressing the heat-activated cation channel TrpA1 cation channel to activate the neuron ^7,52^. ISN inactivation increased water intake in males, consistent with previously reported results in females (**Figure 6H**) ^52^. Inhibiting the ISNs decreased thirsty water seeking in both males and females (**Figure 6I**, **J**). Moreover, activation of the TrpA1 increased water seeking (**Figure 6K**). Thus, the ISNs promote thirsty water seeking and inhibit water intake in both males and females, suggesting that their role in thirst is more complex than simply serving as the initiator of the state of thirst.

## Discussion

We uncovered a male-specific neural circuit for thirsty water seeking, and sex-independent neurons, that together form a basis for understanding the neural representation of appetitive thirst in Drosophila. The male-specific neural pathway progresses from the sex-isomorphic Janu-AstA neuron to the male-specific NPF-M neurons, that converge onto to the mushroom body circuitry functioning sex-independently in thirsty seeking. Key features of this appetitive thirst circuitry include segregated encoding of distinct behaviors by individual circuit elements, multipeptidergic broadcast signaling of NPF-M neurons coupled with cholinergic transmission, and a partial separation of the circuitry for thirst and thirst-dependent learning and memory. Our findings form a basis for deducing how thirst is represented at the single neuron level and how behavioral functions overlap and interact. We predict that NPF-Ms provide a male-specific modification of internal state motivation encoding to influence the interpretation of sensory processing. The male-specific thirst circuit may accommodate male metabolic rate, sustained effort for male mating behavior, or sex-specific behavior in the wild. We also uncovered sex-independent thirsty seeking neurons, including the ISN osmosensors and the OviIN that is thought to form a hub for integrating brain state information.

### Janu-AstA and NPF-M function in thirst

The behavioral roles of Janu-AstA include the promotion of thirsty water seeking, positive valuation, and the inhibition of food foraging, without controlling water intake ^16^. Here we found that Janu-AstA does not regulate either male courtship or female egg-laying, suggesting AstA release from Janu-AstA may be specific to the appetitive phase of ingestive behaviors. The neurons synaptically connected to Janu-AstA reveal possible neural pathways for processing a variety of sensory information into a sex-influenced behavioral drive. The presynaptic source of the thirst signal transmitted by Janu-AstA is not yet known. Input to Janu-AstA is primarily from ascending neurons with strong connectivity to currently undefined sensory inputs. A single Janu-AstA input, the DNp48 descending neuron of unknown function, is potentially sexually dimorphic and therefore may provide male-specific information to Janu-AstA ^39^. DNp48 is downstream to the ISNs via an intermediary neuron named Cowboy, providing a possible path from internal state sensing to Janu-AstA ^19^. Alternatively, NPF-M neurons, serving as a Janu-AstA output, may confer male behavioral specificity to Janu-AstA. Janu-AstA to NPF-Ms connectivity is supported by transsynaptic tracing, the requirement of the AstA receptor AstA-R2 in NPF-Ms for thirsty water seeking, and the male-specificity of AstA release from Janu-AstA for thirsty water seeking. We suggest that thirst sensory information from either ascending neurons and/or the ISNs is contextualized with the help Janu-AstA into a water seeking signal for the NPF-M neurons.

What is the role of the NPF-M in thirsty water seeking? NPF-M neurons are critical for multiple behaviors that are driven by internal state, including thirsty water seeking, thirsty water intake, hungry foraging for food, and male courtship. When NPF-M activity is blocked, all these behaviors are increased, indicating that synaptic release from NPF-M neurons is behaviorally inhibitory. For thirsty water seeking, NPF-M release of acetylcholine is also behaviorally inhibitory, indicating that fast neurotransmission can drive this role. Release of the neuropeptides sNPF, NPF, and DH31 from NPF-Ms promote thirsty water seeking, the opposite effect of acetylcholine. Co-transmission of neuropeptides and neurotransmitters from individual neurons can drive both fast and more persistent effects in downstream circuits, and alter local circuit function more broadly than synaptic connectivity by itself ^53^. Selective depletion of sNPF by thirst suggests that sNPF is responsible for NPF-M peptidergic regulation of thirst behaviors. NPF and DH31 may be modulated at levels below the detection limit or selectively released during water seeking. Alternatively, NPF and DH31 may regulate the development or maintenance of NPF-M circuitry to favor thirsty water seeking behavior.

### Coregulation of courtship and thirst by NPF-M neurons

Although thirst and courtship both require NPF-M neurons, these behaviors appear to use the neurons separably. First, the NPF-M presynaptic P1a neurons that drive courtship are not required for thirsty water seeking. Second, the NPF-M presynaptic Janu-AstA neurons that control thirsty water seeking do not control courtship. Third, the NPF-M neurons only regulate thirsty water seeking when flies are water deprived. Fourth, there exists opposite behavioral roles for NPF release from NPF-M neurons for thirsty water seeking vs. courtship: NPF-M synaptic release inhibits courtship and thirsty water seeking, whereas NPF release from NPF-M neurons inhibits courtship but promotes thirsty water seeking. The role of NPF-M neurons in each motivated behavior may be determined in the context of the predominant internal state, that we predict is encoded elsewhere.

Since the individual NPF-M neurons, SMP712m and ICL010m, have distinct synaptic connectivity and morphology, they may also be distinct behaviorally. Courtship drive is transmitted from P1/P1a neurons that include the pC1x_a-d neurons in the mCNS. The pC1x_b and pC1x_d neurons connect solely to SMP712m whereas pC1x_a connects to both SMP712m and ICL010m. Moreover, ICL010m is presynaptic to the OviIN thirst neuron, the only synaptically connected NPF-M output neuron currently known to regulate thirsty water seeking. Janu-AstA is presynaptic to 2-3 NPF-M neurons, and sNPF, NPF, and DH31 are expressed in all NPF-M neurons, suggesting that thirsty water seeking may be transmitted through the NPF-M neurons as a group.

All defined behavioral roles for the NPF-M neurons can be considered goal-directed behaviors that feature common ongoing types of decisions, but differ by which sensory cues are salient to their ultimate goals. Walking towards a female fly or up a humidity gradient requires locomotion and continuous correction of steering. Both may require continuous prediction of the remaining effort needed to obtain the goal. Both also involve the evaluation of potentially competing internal states and external sensory cues. The synaptic connectivity of NPF-M neurons suggests that sex-specific signals and other information is transformed through the NPF-M neurons to mushroom body, fan-shaped body, and other sex-independent brain structures, creating a sex-biased influence on appetitive behaviors.

### Function of mushroom body circuitry in thirsty water seeking

The mushroom bodies, in addition to their central role in learning and memory, regulate ongoing behavior by integrating information from sensory inputs, internal states, and motor feedback ^54–56^. Our limited survey of neuropeptide receptor and synaptic release requirements suggests that water seeking is broadly distributed into the mushroom body circuitry, including involvement of aversive DANs, appetitive DANs, and the KCs. Thirsty water seeking is similar to hungry food foraging in its broad use of mushroom body circuitry ^57^. The similarities to hungry food foraging extend to requiring the receptors for sNPF and NPF in this circuitry. The physiological response of the mushroom bodies to humidity is not yet comprehensively documented. Water presentation to the fly proboscis, a procedure that provides both humidity and ingestive signals, causes thirst-dependent activation of PAM-β’1 and PAM-γ4 dopamine neurons ^26,44^. The mushroom body KC-γ neurons are third-order with respect to humidity and water taste sensory neurons, where VP1m+VP5_ilPN second order humidity projection neurons activate the KC-γ in response to humidity in thirsty flies ^44,58,59^. Synaptic release from other mushroom body KC neurons, KC-αβ, is required for humidity attraction. However, the humidity attraction behavioral role of the KC-γ and other KC neuron classes has not been tested ^15^. Moreover, the mushroom body circuitry appears to be sex-independent for thirst responses: the aforementioned research tested either females or mixed sex populations, whereas we tested males and/or females ^15,26,44^. Thus, male-specific thirsty seeking is transmitted from the NPF-M to mushroom body circuitry to regulate appetitive seeking behavior.

### Relationship of ongoing and learned thirst behavior in the mushroom body circuitry

The mushroom body neural circuitry also functions in water reward learning and memory. While there may be interactions between ongoing behavior and learned behavior for thirst in the mushroom bodies, our current evidence suggests at least a partial separation. Both PPL1 and PAM dopamine neurons are implicated in water reward learning and memory, including the PPL1-γ1pedc, PPL1-γ2α’1, PAM-β’1, PAM-β’2a, and PAM-γ4 ^25,26,44^. Our findings indicate the PPL1 and PAM neurons also regulate thirsty water seeking, but the specific dopamine neurons responsible are yet to be uncovered for ongoing thirsty water seeking. The PAM-γ4 dopamine neurons, although highly connected to the NPF-M neurons, were dispensable for thirsty water seeking. The PAM-γ4 neurons are activated by water taste or ingestion but not humidity, and they are thought to erase short-term hygrosensory memories that can be formed in thirsty flies, mirroring the transition from appetitive to consummatory behavior ^44^. Neuropeptide signaling to the mushroom body circuitry is important for water reward memory, requiring the neuropeptide receptors NPFR in the PAM-β’2a dopamine neurons and the leucokinin receptor LKR in the PAM-β’2a and PPL1-γ2α’1 dopamine neurons ^25^. Thirsty water seeking requires the receptors for sNPF, NPF, and DH31 in the PAM dopamine neurons, and our partial survey indicates some of these receptors are also required in the PPL1 dopamine neurons.

The mushroom body Kenyon cells are also required for water seeking and water reward memory, but these roles appear to be temporally separable. KC-αβ, KC-α’β’, and KC-γ mushroom body neurons all receive humidity inputs from the VP1m class (Ir68a humidity attraction) and DP1m_adPN (Ir8a humidity response modulation) second order projection neurons ^9,58^. The KC-γ neurons that are activated by humidity and the KC-αβ neurons that are required for thirst-dependent humidity attraction are dispensable for water reward memory acquisition, but they both are required for memory retrieval that occurs 24 hours later ^26^. This raises the possibility that water reward retrieval is triggered by humidity detection when thirsty. We confirmed the requirement for the Kenyon cells in thirsty water seeking and further found that the sNPF receptor is required in these neurons.

### Circuit relationship between male courtship and thirsty water seeking in the mushroom bodies

Male courtship behavior requires a circuit extending from the courtship and aggression P1/P1a neurons to the mushroom body circuitry. PAM-γ4 and PAM-γ5 (collectively known as aSP13) dopamine neurons change their calcium response magnitude in response to the courtship pheromone 11-cis-vaccenyl acetate (cVA) ^60–62^. Moreover, PAM neuron responsiveness to courtship cues is regulated by the P1/P1a neurons. Courtship drive requires the D1-like dopamine receptor Dop1R1 in KC-γ and KC-αβ neurons, and blocking KC synaptic output delays courtship initiation and decreases courtship activity ^63,64^. The PPL1 aversive dopamine neurons also promote male mating behavior ^65^. Neuropeptidergic input to the mushroom bodies for courtship may involve NPF release by NPF-M neurons, but this aspect of courtship regulation remains relatively unexplored ^36^. Thus, ongoing male courtship, like thirsty water seeking and hungry food foraging, is widely distributed in the mushroom body circuitry, suggesting a potential common underlying functional role of the mushroom body across appetitive behaviors.

### Non-sexually dimorphic thirst circuitry

Integration of the OviIN with Janu-AstA suggests some shared function for thirsty seeking. The OviIN, that promotes thirsty water seeking in both males and females, is proposed to be a hub neuron that receives and integrates complex information from groups of functionally related neurons, primarily from neurons centered in the superior protocerebrum ^66^. Janu-AstA falls into a less common category of OviIN input that transmits information from the SEZ region of the brain to the SMP, where Janu-AstA specifically makes synapses in the posterior dorsomedial aspect of the OviIN. We suggest that Janu-AstA provides sensory information about the ongoing thirst state to the OviIN to bias its function. Since Janu-AstA is male-specific for thirsty water seeking while the OviIN isn’t, the OviIN must also gain thirst information from another, as yet unidentified source.

The ISNs are potentially functionally connected to the OviIN via the GNG323 projection neuron. The ISNs inhibit water intake with the insulin-like neuropeptide dILP3, so nonsynaptic communication between these sex-independent thirsty water seeking neurons is also possible ^19^. Our data suggests that ISNs set the state of thirst specifically for ingestion in both sexes, since water seeking is driven by ISN activation and water intake is driven by ISN inhibition, and high osmolarity inhibits the ISNs ^7^. There are multiple thirst sensors in the mouse brain, including osmosensitive and sodium sensitive groups of neurons in the organum vasculosom of the lamina terminalis (OVLT) and the subfornical organ (SFO) nuclei of the lamina terminalis ^3^. These sensors are interconnected to regulate neurons in the median preoptic nucleus (MnPO) that then broadcasts thirst to multiple brain regions. Activation of the ISNs also increases feeding, via the glucagon-like hunger signal adipokinetic hormone (AKH). Thus, ISN activation may initiate prandial thirst associated with feeding, as documented in other species ^2^.

A potential additional thirst pathway may exist for NPF signaling to the mushroom body circuitry, that may or may not be sexually dimorphic for thirsty water seeking. We showed that the NPF receptor NPFR promotes thirsty seeking in the PPL1 dopamine neurons. The PPL1 DANs suppress sugar reward long-term memory via NPF release from the sexually isomorphic L1-l (also known as DAL2) neurons ^67^. The PPL1-α3, PPL1-γ2α’1, and PPL1-γ1pedc require receptors for NPF and sNPF for hungry food foraging ^57^. The L1-l neurons increase activity with thirst and regulate thirsty water seeking ^16,18^. Thus, thirst may activate the L1-l NPF neurons to regulate PPL1 activity and thirsty seeking.

### Why does a male-specific water seeking circuit exist?

Sex differences in the brain and not the body set thirst intensity. Why would males need their own water seeking neural circuitry? Perhaps male-specific behaviors require enhanced thirst sensitivity, for example for the water loss that accompanies courtship song due to spiracle opening and gas exchange ^68^. Courtship opportunity for males may be associated with the presence of a water source in the wild. Drosophila females tend to stay on energy rich food sources that also contain water, whereas males tend to disperse ^69^. Seeking water, food, and mates may thus be coupled behaviorally. Finally, the NPF-M pathway may translate male-specific energetic needs onto sexually isomorphic brain functions. NPF-M-dependent coupling could drive temporal matching to the differences in metabolic rate in male flies ^70^.

## Methods

### Fly strains and culturing conditions

*Drosophila melanogaster* strains, antibodies, and key reagents are listed in **Table S2**. *Drosophila* strains were outcrossed for five to eight generations to the Berlin genetic background, except for *fru* mutants and the *NPF-Gal4;fru-FLP* transgenic strain that were in the Canton-S genetic background. Flies were raised on standard food containing agar (1.2% w/v), cornmeal (6.75% w/v), molasses (9% v/v), and yeast (1.7% w/v) at 25°C and 60% humidity in a 12:12 light:dark cycle in incubators from Darwin Chambers. For experiments with *UAS-TrpA1*, flies were reared at 21°C prior to testing. For *NPF-M-spGal4>UAS-RNAi*, adult F1 progeny were held at 29°C and 60-70% humidity for 3-4 days prior to testing to enhance transgene activity. For experiments with *UAS-STOP-effector/FLP*, flies were aged 1-3 additional days to enhance Flp-out of the STOP cassette. For experiments with *trans*-Tango, flies were aged to 10-20 days old at 25°C before dissection to maximize signal intensity ^35^. Male mating status was manipulated as follows. To generate sexually experienced males, 10 males were paired with 30 genetic background females at 1-2 days old and flipped into fresh cultures every day for 3 days. Female flies were removed from the vial through brief CO_2_ anesthetization the day prior to experimentation. To generate sexually inexperienced males, groups of 20 males were flipped into fresh cultures daily for 3-4 days and split into two groups of 10 through brief CO_2_ anesthetization the day prior to experimentation.

### NPF-M identification in single cell RNA sequencing datasets

Preexisting single cell RNAseq data was manually analyzed to identify *NPF*-positive *fru*-positive (male-specific) cell bodies as NPF-M neurons ^42,43^. *Split-Gal4* activation domain transgenes derived from enhancer regions of genes with enriched expression in these neurons were screened against *NPF-T2A-DBD* to identify *NPF-M-spGal4* (*VT002860-p65.AD*) ^67^.

### Water and food deprivation

For water deprivation, flies were placed in empty culture vials containing 1 mL 5% sucrose (S5-500, Thermo Fisher) that was dried onto a 2.5 × 15 cm strip of Whatman 3 MM filter paper (WHA3030662, Sigma-Aldrich). For food deprivation, flies were placed into empty culture vials containing 3 MM filter paper soaked with 800 μL of UltraPure H_2_O (10977-023, Thermo Fisher).

### Water seeking and food seeking

Groups of approximately 20 3-6 day old adult males were collected 1–2 days prior to the experiment. One group is an n = 1. Sources of water (with 0.2% w/v erioglaucine, 229730050, Thermo Fisher) were presented to flies in the cap of a 0.65 mL microcentrifuge tube. Water was made inaccessible by glueing (698, DAP) a 300 μm mesh grid (U-CMN-300-A, Component Supply) onto the cap. Dry, rough cut cane sugar (Pure Cane Rough Cut Cubes, La Perruche) was presented in the cap of a 1.5 mL microcentrifuge tube for food seeking. Thin-walled Plexiglass behavioral chambers were designed as two side-by-side arenas with each arena measuring 45×75×10 mm. Chambers were designed and built by IO Rodeo (Pasadena, CA). The chambers and water/food sources were acclimated to the testing temperature in a Peltier incubator (IN45, Torrey Pines Scientific), and flies were acclimated into the chambers for 10 min prior to introducing the water/food source. Humidity, measured using an EK-H4 multiplexer with SHT71 sensors (Sensirion), was maintained at 30–40% relative humidity. Flies were filmed from above with Logitech C510 web cameras at 30 fps with the arena placed on white light LED panel (Edmund Optics). The number of flies on the water/food source was divided by total number of flies to determine the fraction of flies on the water/food source.

### Water intake

Flies were glued to microscope slides with clear nail polish while cold anesthetized and then allowed to recover at 25°C and 60% humidity for 1-2 hours. Water consumption time was measured by repeatedly presenting the proboscis with a water drop extruded from the end of a 5 μL calibrated micropipette (53432-706, VWR), until flies failed to extend the proboscis to four sequential presentations. Behavior was recorded on a stereoscope with an AmScope MD500 microscope eyepiece camera at 30 fps at 640 x 480. 0.2% w/v erioglaucine (229730050, Thermo Fisher) was added to increase contrast. The first bout length was the time from first engagement to first complete disengagement with the water drop. Pumping rate was determined by manual counts of cibarial contractions through video recordings during the initial 10 s of the first engagement, or the entire first bout length if a fly disengaged earlier. If the fly never drank from the water drop, the data was excluded from analysis.

### Courtship behavior

Courtship assessment was performed in an upside-down 24-well plate. One male and one virgin female fly were gently introduced to a well that was then capped with a cut bottle plug (59-194P, Genesee Scientific), and their activity was recorded from above for 10 min in sets of 4 mating pairs across 4 wells. Courtship was recorded with a JVU250 camera at 30 fps while the 24-well plate sits atop a white light LED panel set to the lowest illumination setting. For experienced males, 4-6 day old males were collected and stored with virgin females in a 1:1 ratio for 2-3 days. For inexperienced males, males were collected and isolated as pupae to eclose in a socially isolated environment. The males were socially isolated for 3 days before being tested in the courtship assay. Courtship index was calculated as ratio of the time spent courting over the 10-min session. Male chasing behavior, horizontal wing extension, and copulation were counted as courting behavior.

### Egg-laying behavior

For mated female egg-laying behavior, individual mated females were placed into vials with a single genetic background male and flipped every day onto fresh food for 4 days. The eggs from each vial were counted and the number of eggs laid each day was averaged. For virgin female egg-laying behavior, virgin females were collected, distributed into groups of 4-5 virgin female flies per vial, and flipped every day for 5 days. Eggs were counted from each vial and added together.

### Water deprivation lethality

For *R13F02>TeTx* water deprivation lethality experiments, 20 male flies were placed into dry sucrose vials and the number of living flies was counted every 0.5-1 hr. For male:male vs. male:female water deprivation lethality experiments, 20 sex-deprived male flies were placed into dry sucrose vials with either 10 other males or 30 mated females, and the number of dead males was counted hourly.

### Immunohistochemistry

Adult fly brains were dissected in 1x PBS (BP399, Thermo Fisher) with 0.05% Triton-X 100 (X100-500ML, Sigma-Aldrich) (0.05% PBT), fixed overnight at 4°C in 0.05% PBT with 2% paraformaldehyde (50-980-493, Thermo Fisher) in 0.5mL Protein LoBind tubes (13-698-793, Thermo Fisher). Brains were then washed with 1x PBS with 0.1% Triton-X 100 (0.1% PBT) for 20 min at room temperature 5 times, blocked in 0.5% PBT with 5% normal goat serum (733250, Lampire Biological laboratories) and 0.5% bovine serum albumin (A7906-50G, Sigma-Aldrich) for 2 hr at room temperature, and incubated with primary antibody diluted in block for 1-3 days. Brains were washed with 0.1% PBT for 20 min at room temperature 5 times, incubated in secondary antibody diluted in 0.1% PBT for 1-2 days, washed with 0.1% PBT for 20 min at room temperature 5 times, and incubated in Vectashield or Vectashield PLUS (H-1000 or H-1900, Vector Laboratories) overnight at 4°C. Samples were mounted in Vectashield and imaged on a Zeiss LSM-880 confocal microscope with a 20x, 40x, or 63x objective. Image stacks were processed in Fiji, and brightness and contrast were adjusted in Adobe Photoshop. For intestines, the tissue was incubated in Vectashield PLUS with DAPI (H-2000, Vector Laboratories). For MultiColor FlpOut experiments, flies were heat shocked in a 37°C water bath for 40-45 min and aged for 2 days at 25°C and 60% humidity. Foreleg and wing imaging was done directly on freshly dissected tissues in PBT. To determine neuropeptide levels with thirst, wild-type genetic background brains were immunostained as described above. All brains were placed onto the same slide and were imaged on the same day, using fixed confocal settings with a 20x objective. Z-stacks were taken at 1.5 μm interval and to the minimum z-depth that captured all NPF-M cell bodies, so that all stacks were the same depth. These were maximum intensity z-compressed. The mean intensity was measured for a 22 μm square area encompassing single NPF-M cell bodies and for a background area immediately adjacent, using Fiji. The two most strongly staining cell bodies for each hemisphere were recorded. The values reported are signal minus background, expressed as the fraction of the maximum possible signal.

### Connectome analysis

The three-dimensional closest distance between two neurons in the mCNS connectome was determined with custom R code. Neuron meshes were obtained from the male CNS dataset (v0.9), converted from nm to μm, and cleaned up using vcgclean from the Rvcg package. A k-dimensional tree was used to find the closest vertex between pairs of neurons. Janu-AstA neurons (mCNS 14134 and 524224) were compared to ISNs (131216, 179881, 524875, and 184281026) and NPF-M neurons (524998, 20875, 56766, and 21678).

## Statistics and data analysis

Genotype and treatment were blinded for all behavioral counts. Statistics were calculated with Prism 10 (GraphPad). Statistical tests used are listed per Figure. For experiments with one experimental group and multiple control groups, a One-way ANOVA with a Holm-Šídák’s multiple comparisons test was performed. If the data failed the Brown-Forsythe test, a Brown-Forsythe test with Dunnett’s T3 multiple comparisons test was performed instead. If the data failed the Shapiro-Wilks test regardless of the Brown-Forsythe test, a Kruskal-Wallis test with Dunn’s multiple comparison test was used. For experiments with two groups, a two-tailed t test was used. If the data had unequal standard deviations, Welch’s t test was used. If the data failed the F-test, Mann-Whitney t test was used. Error bars are the standard error of the mean (SEM). Dots overlaid on bar graphs are the value measured for individual n. ****p≤0.0001; ***p≤0.001; **p≤0.01; *p≤0.05; and ns, p>0.05. Complete statistical analyses are included in **Table S3**.

## Data availability

All data is included in the manuscript and in **Data S1**.

## Code availability

All computer code is available via GitHub at https://github.com/flowderf/Sexually-Dimorphic-Thirsty-Water-Seeking.

## Author Contributions

B.P.W.: conceived of the study, performed most of the experiments, contributed to methodology, contributed to study design, wrote and edited the manuscript. X.Z., S.S., G.K., N.F.: performed experiments. F.W.W. led the study, performed experiments, contributed to methodology, contributed to study design, created figures, wrote and edited the manuscript.

---

**Figure S1.**
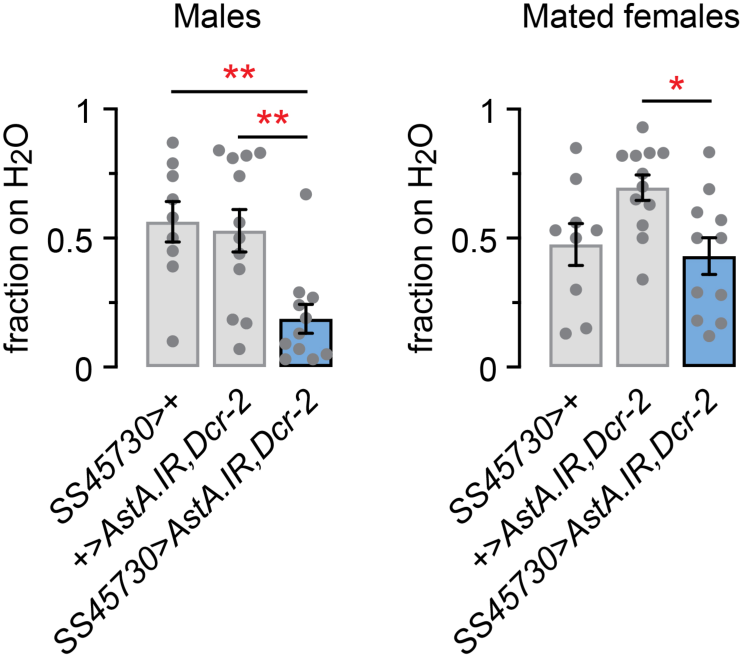
Water seeking role of AstA tested with a highly specific Janu-AstA split-Gal4. RNAi reduction of AstA in Janu-AstA with SS45730-spGal4 reduced water seeking in males (left), but not females (right). Left and right: one-way ANOVA/Holm-Šídák’s.

**Figure S2.**
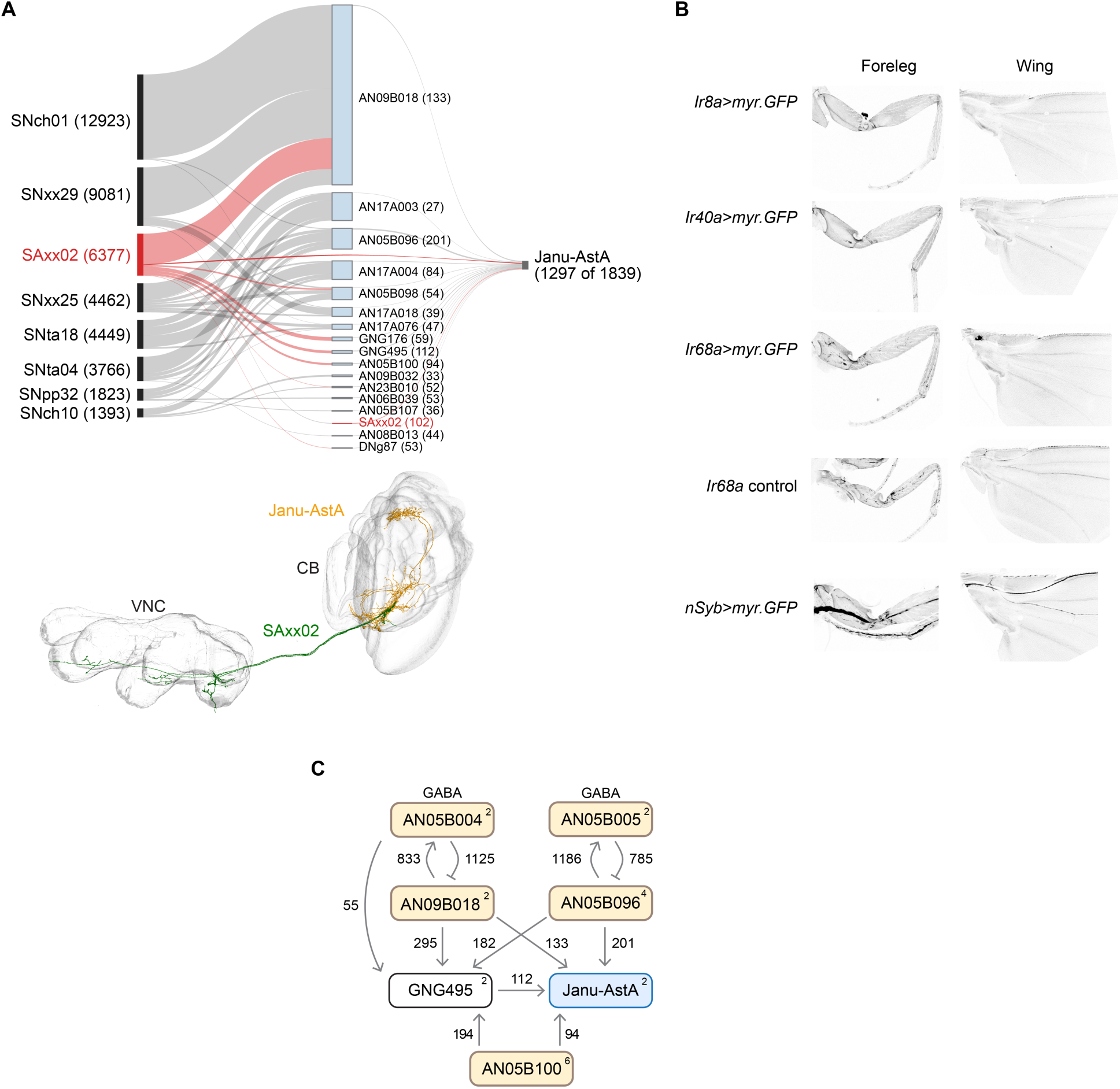
Janu-AstA synaptic input connectivity. **A**. Sensory and ascending neuron inputs to Janu-AstA, organized by sensory neuron type (left). Ascending neurons are the middle layer. Numbers indicate synapses made to the next layer. SNxx29 is presynaptic to the same group SAxx02 neurons that directly connect to Janu-AstA; this SAxx02 connection to Janu-AstA is represented redundantly. Below: SAxx02 neurons that are directly presynaptic to Janu-AstA in the mCNS male connectome. **B**. Lack of expression of hygrosensory receptors Ir8a, Ir40a, and Ir68a in the foreleg or the wing. *nSyb>myr.GFP* labels major nerve tracts in both appendages as a positive control. **C**. Diagram of symmetric ascending neuron input to Janu-AstA and GNG495.

**Figure S3.**
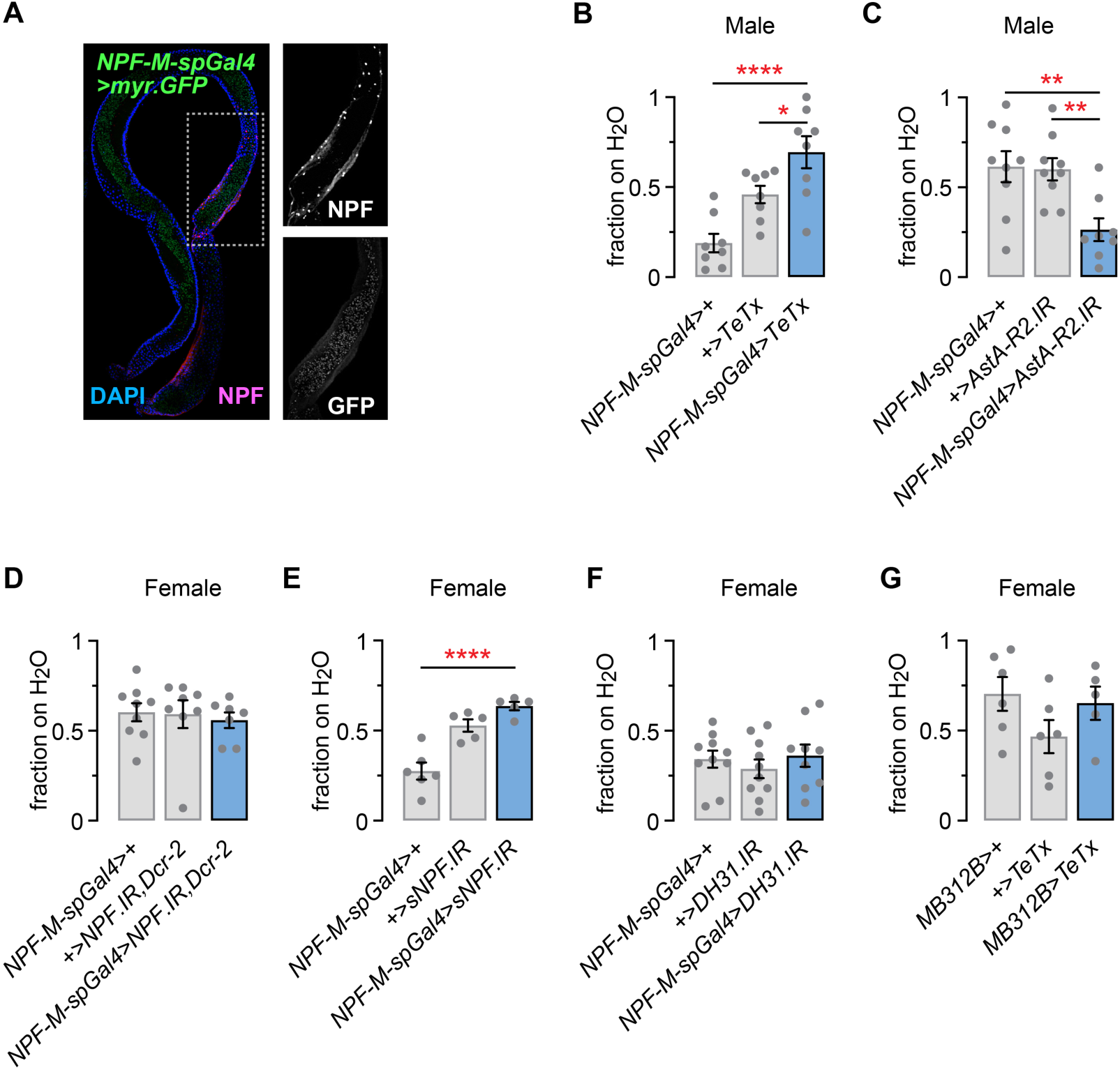
*NPF-M-spGal4* recapitulates *NPF-Gal4,fru-FLP* as a driver for the NPF-Ms and neuropeptide RNAi effects on female water seeking. **A**. Adult male Drosophila intestine stained for GFP (green), NPF (magenta), and DAPI (blue). Green signal in the intestinal lumen is fluorescence from gut microbiota. Right: NPF and GFP channels from dashed box region. **B**. Synaptic inactivation of NPF-M neurons with *NPF-M-spGal4* increases thirsty water seeking in males. One-way ANOVA/Holm-Šídák’s. **C**. RNAi reduction of AstA-R2 in NPF-Ms with *NPF-M-spGal4* reduces thirsty water seeking in males. One-way ANOVA/Holm-Šídák’s. **D**. RNAi reduction of NPF with *NPF-M-spGal4* has no effect on thirsty water seeking in females. One-way ANOVA. **E**. RNAi reduction of sNPF with *NPF-M-spGal4* has no effect on thirsty water seeking in females. One-way ANOVA/Holm-Šídák’s. **F**. RNAi reduction of DH31 with *NPF-M-spGal4* has no effect on thirsty water seeking in females. One-way ANOVA.

**Figure S4.**
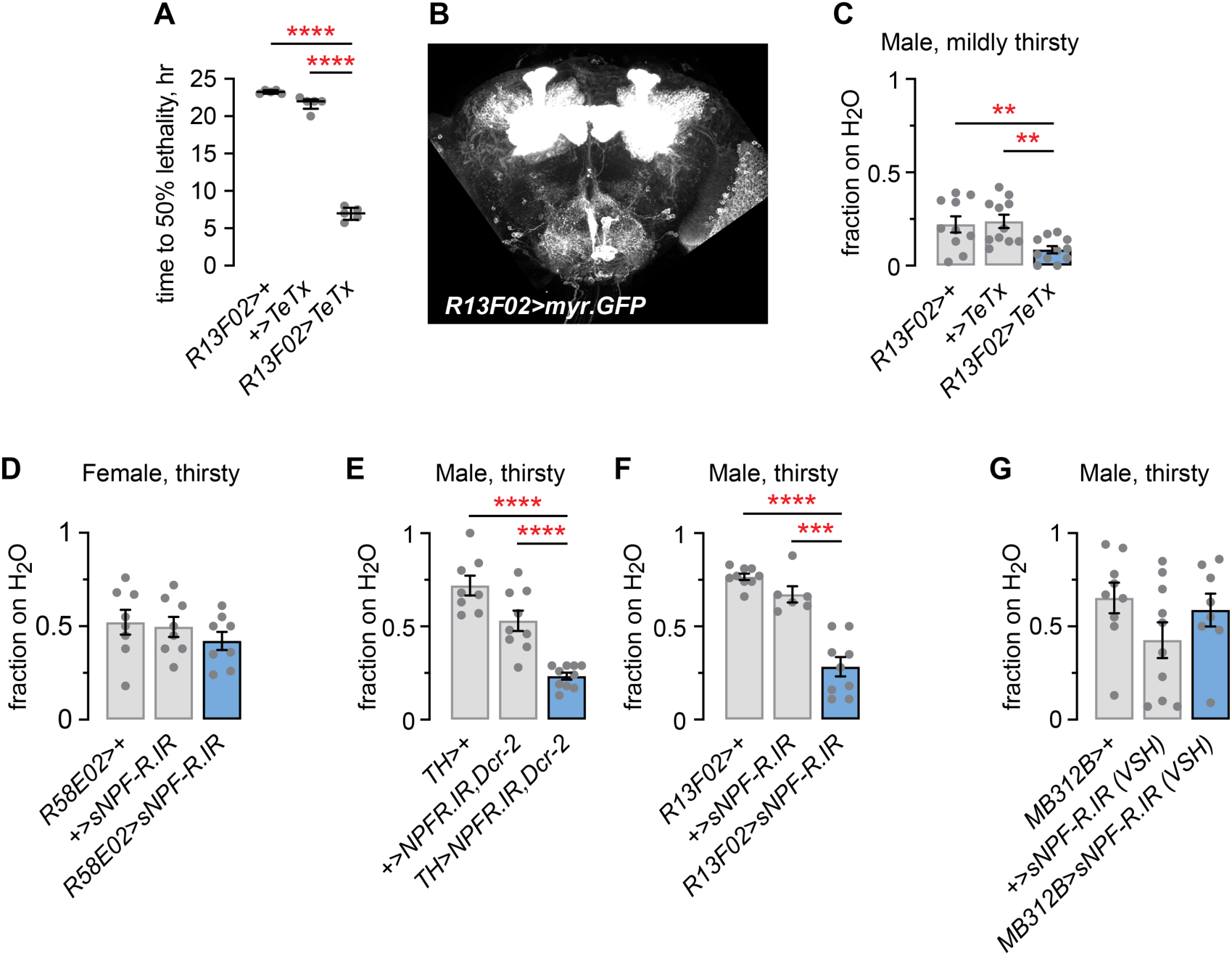
Role of mushroom body circuitry in thirsty water seeking. **A**. Time to 50% lethality for *R13F02>TeTx* males when water deprived on dry sucrose. One-way ANOVA/Holm-Šídák’s. **B**. Z-compressed confocal image of *R13F02-Gal4>myr.GFP*, overexposed to reveal neurons in addition to the mushroom body Kenyon cells. **C**. Water seeking in *R13F02>TeTx* flies that were briefly water deprived (4 hr). One-way ANOVA/Holm-Šídák’s. **D**. RNAi reduction of sNPF-R in *R58E02-Gal4* appetitive DANs, no effect on female thirsty water seeking. One-way ANOVA. **E**. RNAi reduction of NPFR in aversive dopamine neurons with *TH-Gal4* decreases thirsty water seeking. One-way ANOVA/Holm-Šídák’s. **F**. RNAi reduction of sNPF-R in mushroom body Kenyon cells decreases thirsty water seeking. Brown-Forsythe/Dunnett’s T3. **G**. A second sNPF-R RNAi, *VSH330347*, did not alter thirsty water seeking when expressed in the PAM-γ4 neurons in males. One-way ANOVA.

**Figure S5.**
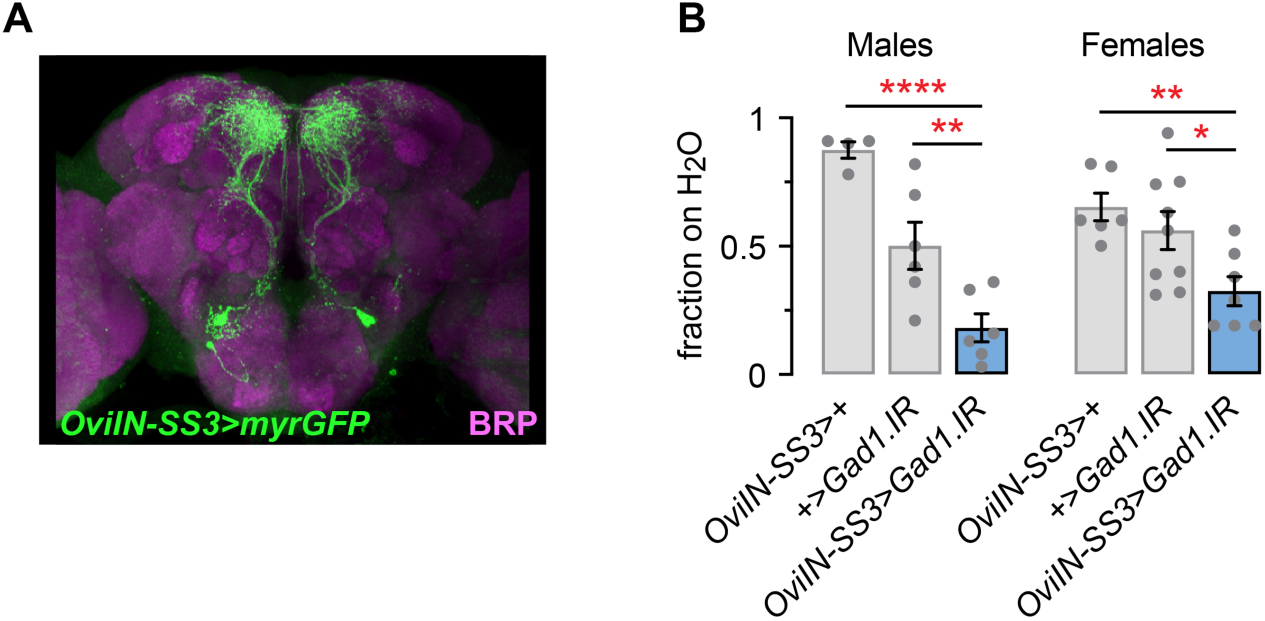
Recapitulation of the OviIN thirsty seeking behavioral phenotypes with a second OviIN *split-Gal4*. **A**. Expression pattern of *OviIN-SS3 split-Gal4*. **B**. RNAi reduction of *Gad1* in either males or females decreases thirsty water seeking. Left and right: one-way ANOVA/Holm-Šídák’s.

